# Are single peripheral measurements of baseline oxytocin in saliva and plasma reliable biomarkers of the physiology of the oxytocin system in humans?

**DOI:** 10.1101/2020.07.14.202622

**Authors:** Daniel Martins, Anthony Gabay, Mitul A. Mehta, Yannis Paloyelis

**Author notes:** **Corresponding author**: Yannis Paloyelis, PhD, Department of Neuroimaging, Institute of Psychiatry, Psychology and Neuroscience, King’s College London, De Crespigny Park, London SE5 8AF, United Kingdom. **Category of manuscript** Archival report.

## Abstract

**Background:** Single measurements of salivary and plasmatic oxytocin are used as indicators of the physiology of the oxytocin system. However, questions remain about whether they are sufficiently stable to provide valid biomarkers of the physiology of the oxytocin system, and whether salivary oxytocin can accurately index its plasmatic concentrations.

**Methods:** Using radioimmunoassay, we measured baseline plasmatic and/or salivary oxytocin from two independent datasets. Dataset A comprised 17 healthy men sampled on four occasions approximately at weekly intervals. We administered exogenous oxytocin intravenously and intranasally in a triple dummy, within-subject, placebo-controlled design and compared baseline levels and the effects of routes of administration. Dataset B comprised baseline plasmatic oxytocin measurements from 20 healthy men sampled on two separate occasions. Additionally, in dataset A, we tested whether salivary oxytocin can predict plasmatic oxytocin at baseline and after intranasal and intravenous oxytocin administration.

**Results:** Single measurements of plasmatic and salivary oxytocin showed poor reliability across visits in both datasets. Intranasal administration of exogenous oxytocin increases salivary oxytocin, but intravenous administration of a considerable dose does not produce any changes. Saliva and plasma oxytocin did not correlate at baseline or after administration of exogenous oxytocin.

**Conclusions:** Our findings question the use of single measurements of baseline oxytocin concentrations in saliva and plasma as valid biomarkers of the physiology of the oxytocin system in humans. Salivary oxytocin is a weak surrogate for plasmatic oxytocin. The increases in salivary oxytocin observed after intranasal oxytocin most likely reflect unabsorbed peptide and should not be used to predict treatment effects.

## Introduction

In the last two decades, a wave of studies have sought to investigate the role of oxytocin in human normal and impaired socio-affective behaviour and cognition(1, 2). Several methodological approaches have been adopted to this effect. Predominantly these include: i) the measurement of psychological or neurobiological outcomes after the intranasal administration of exogenous oxytocin compared to placebo(3); ii) the assessment of associations between single measurements of the concentration of endogenous oxytocin in peripheral fluids (mainly blood and saliva) at rest(4) and individual differences in neurobehavioral phenotypes(5), psychiatric disorder status and/or symptom severity(6).

The latter approach is based on two assumptions. The first is that single measures of baseline levels of endogenous oxytocin in the biological fluids of peripheral compartments can accurately index oxytocin release in the brain(4). The second is that single measures provide reliable estimates of baseline levels of endogenous oxytocin in plasma or saliva. Definitive answers regarding the first assumption, whether it regards baseline or evoked release following some intervention, remain to be obtained(7).

Here, we investigate the second assumption, which is a prerequisite if single measurements of baseline levels of endogenous oxytocin are to be used as a valid biomarker of the physiology of the human oxytocin system(8). Currently, we lack evidence that single measures of endogenous oxytocin in saliva and plasma at rest are stable enough to provide a valid biomarker. Such evidence is urgently required, given reports that plasma and saliva levels of oxytocin are frequently altered during neuropsychiatric illness and that they co-vary with clinical aspects of disease(6).

The measurement of salivary oxytocin has been proposed as a surrogate for blood plasma levels(4). Compared to blood sampling, saliva collection presents several logistical advantages(9). However, the validity of this approach remains unclear. While elevations in plasmatic oxytocin after intranasal administration are likely to result from capillary absorption in the nasal cavity(10), elevations in saliva oxytocin could result from mucociliary clearance of intranasally delivered oxytocin from the nasal cavity to the oropharynx (“drip-down” oxytocin)(11, 12). This question can be illuminated by ascertaining changes in salivary oxytocin following the intravenous administration of oxytocin, eliminating the confound of drip-down oxytocin. In a previous study using enzyme-linked immunosorbent assay, the administration of low and medium doses of intranasal oxytocin (8 and 24 IU) elevated oxytocin concentrations in saliva, but the administration of a low dose (1 IU) intravenously had no effect. Moreover, concentrations of oxytocin in plasma and saliva did not correlate at baseline or after the administration of exogenous oxytocin for either route(13). These data suggest that salivary oxytocin is a weak surrogate measure for peripheral blood levels. However, questions remain about whether the same results would have been achieved by using a more sensitive method of quantification, such as radioimmunoassay (currently the gold-standard for oxytocin quantification(14)), or higher doses of oxytocin administered intravenously.

Here, we aimed to characterize the reliability of both salivary and plasmatic single measures of basal oxytocin in two independent datasets, to gain insight about their putative validity as biomarkers. Additionally, we investigated whether salivary oxytocin concentration reflects plasmatic oxytocin by examining i) if the intravenous administration of exogenous oxytocin increases the concentration of salivary oxytocin; and ii) the correlation between plasmatic and salivary oxytocin levels at baseline and after exogenous oxytocin administration. For all analyses, we followed current gold-standard practices in the field and assayed oxytocin concentrations using radioimmunoassay in extracted samples(14).

## Methods and Materials

### Participants

**Dataset A** included 17 healthy, right-handed, male volunteers (mean age (SD) = 23.75 (5.10); range = 18-34) who contributed samples over four visits, as part of a larger study. **Dataset B** (independent replication study) included 20 healthy, right-handed, male volunteers (mean age (SD) = 24.8 (3.70); range = 21-37), who contributed samples over two visits as part of a different study (both studies described below). All participants had no history of psychiatric disorders or substance abuse, scored negatively on a screening test for recreational drug use, and did not currently use any medication. They were advised to avoid heavy exercise, alcohol or smoking the day before scanning and avoid any drink or food within two hours before scanning. Both studies were approved by King’s College London Research Ethics Committee (Dataset A: PNM/13/14-163; Dataset B: PNM/14/15-32). For both studies, our sample size and number of samples collected per individual would have allowed us to detect intra-class correlation coefficients (ICC) of at least 0.70 (moderate reliability) with 80% of power(15).

### Study design

#### Dataset *A*

Participants were recruited to participate in a double-blind, placebo-control, triple-dummy, cross-over MRI study exploring the effects of various methods of exogenous oxytocin administration on cerebral physiological responses at rest(16). Participants received, in counterbalanced order, over four consecutive visits spaced on average 8.80 days apart (SD 5.72; range 3-28), approximately 40IU of intranasal oxytocin, either with a nasal spray or the SINUS nebulizer (PARI GmbH), 10IU of oxytocin intravenously, or placebo. In each visit, blood samples were collected at baseline, immediately after each treatment administration, and at six time points post-administration, with the last sample acquired when participants came out of the scanner at about 115 minutes post-administration (Fig. 1). Saliva samples were acquired at baseline and together with the last blood sample. For the purposes of this report, we use the plasmatic and salivary oxytocin measurements that were obtained at baseline and at 115 minutes post-administration.

**Fig. 1.**
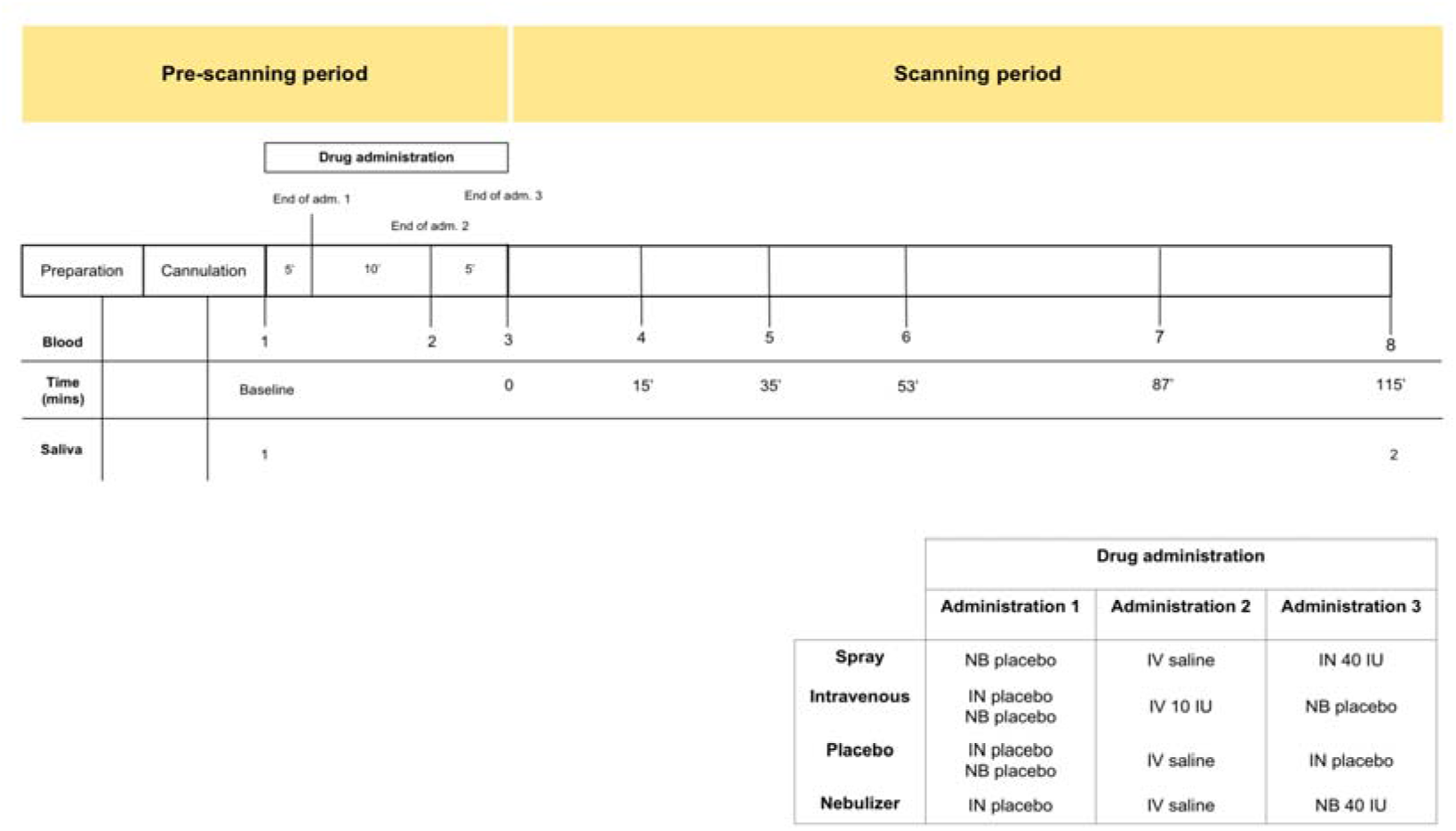
Schematic representation of the design of study A. All subjects received first an administration of intranasal placebo - either by spray or nebuliser, then an intravenous administration of oxytocin (10 IU)/saline and then an intranasal administration of oxytocin (40 IU)/placebo, either by spray or nebuliser. Following drug administration participants were placed in a Magnetic Resonance Imaging scanner for eight resting arterial spinal labelling (ASL) regional blood flow images of the brain and one resting BOLD fMRI scan. Saliva samples were collected before any drug administration (baseline) and after the scanning session (at 115 minutes post-administration of oxytocin). Plasma samples were collected before any drug administration, after any administered drug and then at several time-points during scanning session. For the current study, only time-points where saliva and plasma were concomitantly collected were considered – baseline and after scanning session. Detailed plasmatic pharmacokinetics of each route/method have been presented elsewhere(16). Adm. – administration; mins - minutes.

All visits were identical in structure (duration ~ 3.5 hours). Upon arrival, participants gave consent and completed the required questionnaires. We then fitted an intravenous cannula on each arm of our participants (one for the intravenous administration and another for blood-sampling). Treatment was administered according to the study protocol (Fig. 1). At the end of the treatment administration, participants were taken to an MRI scanner where we obtained a number of resting state and structural scans over the course of the next two hours.

#### Dataset B

Participants were recruited to participate in a study examining the effects of MDMA on social cognition(17). Briefly, baseline blood samples were obtained 15 min before MDMA/placebo administration on two separate occasions, spaced on average 9.30 days apart (SD = 5.70 days; range: 7–31 days).

### Saliva and plasma collections

Blood was collected in ethylenediaminetetraacetic acid vacutainers (Kabe EDTA tubes 078001), placed in iced water and centrifuged at 1300 × g for 10 minutes at 4°C within 20 minutes of collection and then immediately pipetted into Eppendorf vials. Samples were immediately stored −80°C until analysis. Saliva samples were collected using a salivette (Sarstedt 51.1534.500) and then centrifuged and stored in the same manner as blood samples.

### Quantification of oxytocin in plasma and saliva samples

For both datasets, plasma and saliva oxytocin levels were analysed by a third party (RIAgnosis, Munich, Germany) using a Radioimmunoassay (RIA), as previously described(18). Plasma samples were extracted before quantification. Saliva samples were not extracted prior to quantification since unpublished data from *RIAgnosis* found no differences in oxytocin concentrations between extracted and simply evaporated saliva samples. RIA has been previously standardised and validated and represents the gold-standard for oxytocin measurement in biological fluids(18–24). The detection limit is in the 0.5 pg/sample range, depending on the age of the tracer. Cross-reactivity with vasopressin, ring moieties and terminal tripeptides of both oxytocin and vasopressin and a wide variety of peptides comprising 3 (alpha-melanocyte-stimulating hormone) up to 41 (corticotrophin-releasing factor) amino acids are <0.7% throughout. The intra- and inter-assay variabilities are <10%(18).

### Dataset A

#### Reliability analysis

Reliability refers to the reproducibility of values of a measurement in repeated trials on the same individuals(25). Reliability can be quantified using two sets of metrics providing complementary information: absolute and relative reliability. Absolute reliability is the degree to which repeated measurements within the same subject vary over time(25). Relative reliability is the degree to which individuals maintain their position in a sample of subjects measured over time(25). Absolute and relative reliability of plasma and salivary oxytocin measurements were estimated using the within-subject coefficient of variation (CV) and the intra-class correlation coefficient (ICC), respectively. ICC was estimated in a two-way mixed model, single measures, absolute agreement(26). We first estimated the reliability across all four sessions, and subsequently for each pair of visits to assess if time-interval between sample collections may impact on reliability indexes estimation. Only participants presenting baseline measures across all four sessions were included in the reliability analysis, as previously suggested(27).

#### Mean concentrations across visits

Mean concentrations of saliva and plasma oxytocin across the four visits were compared using repeated-measures one-way analysis of variance.

#### Treatment effects

The effect of treatment on blood/saliva oxytocin concentration were assessed using a 4 × 2 repeated-measures two-way analysis of variance Treatment (four levels: Spray, Nebuliser, Intravenous and Placebo) × Time (two levels: Baseline and post-administration). Post-hoc Tukey for multiple comparisons was used to investigate simple effects following a significant interaction.

#### Association between salivary and plasmatic oxytocin levels

We assessed correlations between salivary and plasmatic concentrations of oxytocin sampled at baseline and post-administration. For the baseline measurements, we pooled data across treatment levels since there were no differences between groups on mean baseline concentrations of oxytocin. To account for the non-independence among the four data points within each subject, we used multilevel correlation, where we modelled participant as a random effect. For the post-administration measurements, we calculated Pearson’s correlation coefficient for each treatment level separately because group differences in mean scores on these measures might result in illusory correlations if the drug and placebo samples were pooled together(28). Since our sample was relatively small and therefore the lack of significant correlations between salivary and plasmatic oxytocin could simply reflect lack of sensitivity, we followed this frequentist correlation analysis with Bayesian statistics to quantify relative evidence for both the null and alternative hypotheses.

#### Outliers and missing values

Salivary oxytocin concentrations were missing for three participants, and plasmatic oxytocin concentration for one participant. One measure of baseline oxytocin in saliva and two post-administration measures in the nebuliser condition were discarded after they had been identified as outliers. Outliers were identified using the *outlier labelling rule*(29). A total of 13 and 16 participants were included in the reliability analysis of salivary and plasmatic oxytocin, respectively.

### Dataset B

#### Mean concentrations across visits

Mean concentrations of plasma oxytocin across the two visits were compared using a paired T-test.

#### Reliability analysis

Absolute and relative reliability of plasma oxytocin measurements were analysed for the two baseline measures obtained from each of the two visits, following the methods described for dataset A.

#### Outliers and missing values

There were no missing values. One baseline measure for one of the visits was discarded after being identified as an outlier. A total of 19 participants were included in the reliability analysis.

Increasing the number of observations per individual and averaging across several samples collected on different occasions is an approach commonly used to control within-individual variation and maximize reliability(30). Hence, we expanded our reliability analysis by asking how many additional measures of the same individual would be theoretically required to achieve different levels of reliability of a hypothetical averaged measure, based on the initial reliabilities estimated in the studies A and B. This number was calculated using the Spearman-Brown prediction formula(31). For these calculations, we considered cut-offs of ICC = 0.50 (fair reliability), 0.70 (moderate) and 0.80 (good) as suggested by Koo and colleagues(26).

### Statistical analysis

All statistical analyses were conducted on log-transformed oxytocin concentrations given the deviations of these measurements from a Gaussian distribution. The statistical analysis investigating treatment effects on salivary and plasmatic oxytocin were performed using SPSS (version 24, IBM, Armonk, NY, USA). The frequentist and Bayesian correlations were implemented in the *correlation* package from R (version 3.5.3), using bootstrapping 1000 samples. An increase in Bayes Factor (BF) in our analyses corresponds to an increase in evidence in favour of the null hypothesis (further details in Supplementary). Figures were produced using the *ggplot* package from R (version 3.5.3). P < 0.05 (two-tailed) was set as threshold of statistical significance for all analyses.

## Results

### Baseline salivary and plasmatic oxytocin concentrations across visits

We did not identify any significant differences in mean concentration of baseline oxytocin in saliva or plasma samples across the four visits of dataset A (<u>Plasma:</u> F(1.47, 22.04) = 0.51, p = 0.55; <u>Saliva:</u> F(1.06, 12.82) = 0.88, p=0.38) (Fig. S1). We also did not observe any differences in mean concentrations of baseline oxytocin in plasma across the two visits of dataset B (T(19) = 0.63, p = 0.54). However, in a quick inspection of Fig.2, we can observe that the levels of baseline salivary and plasmatic oxytocin fluctuated considerably from one visit to another in most individuals (Fig. 2).

**Fig. 2.**
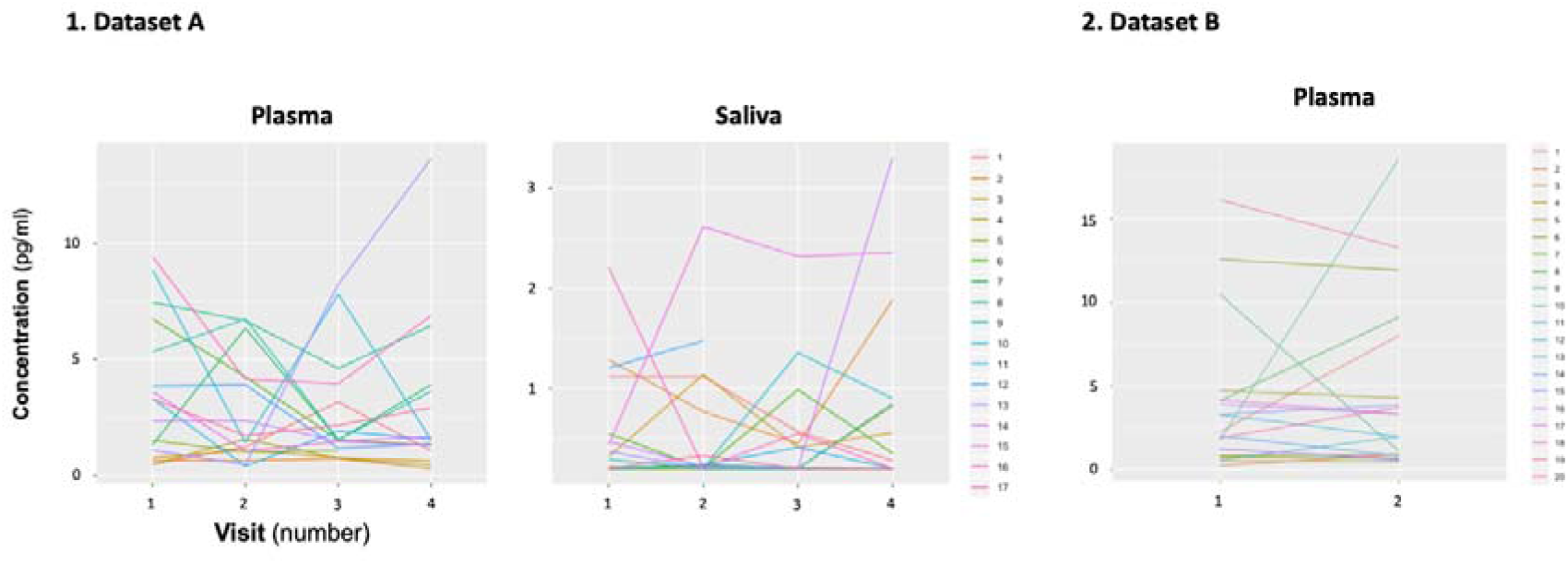
Within-individual variation of baseline measurements of oxytocin in plasma and saliva samples across visits. Baseline plasmatic and salivary oxytocin fluctuate from one visit to another for most individuals (Dataset A). We replicated this trend in an independent dataset for plasma (Dataset B). Each coloured line represents one individual.

### Reliability of single oxytocin measurements in the plasma and saliva

#### Dataset A

##### Reliability analysis across the four visits

We estimated the ICC of single oxytocin measurements in saliva to be 0.23 and in plasma 0.29. The mean CV was 63% for saliva and 57% for plasma measurements. We estimated the number of measurements that would have been required to achieve good reliability (ICC = 0.80) of a putative averaged measure to be 13.39 for saliva and 9.79 for plasma (Table 1).

**Table 1.**
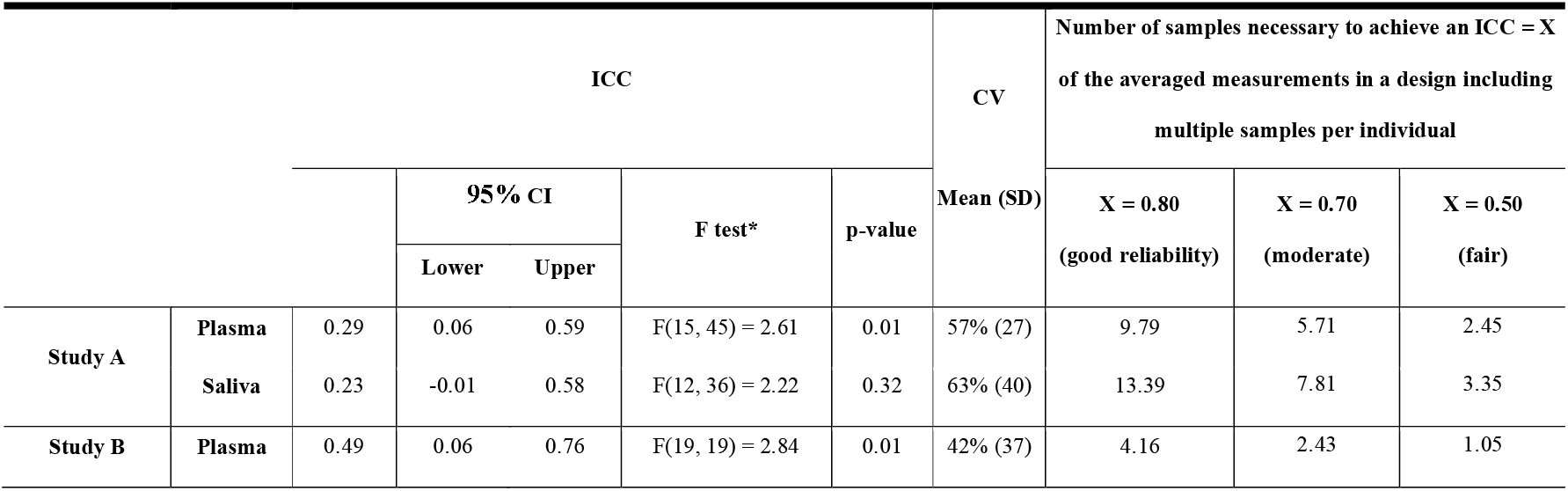
Estimates of absolute and relative reliability of oxytocin measurements in saliva and plasma samples. Absolute and relative reliability were analysed using the within-subject coefficient of variation (CV) and the intra-class correlation coefficient (ICC), respectively. We also present the number of additional measurements of the same individual that would be theoretically required to achieve different levels of reliability (ICC = X) of a hypothetical averaged measure, based on the initial reliabilities estimated for our datasets A and B. This number was calculated using the Spearman-Brown prediction formula. CI – confidence interval; SD – Standard Deviation; *H0: ICC is not significantly different from 0.

##### Reliability analysis for each pair of visits

Detailed descriptions of the ICCs and coefficients of variation estimated for each pair of the four visits included in our analysis of dataset A are presented in Table S1. For plasma, we found higher estimates of ICC for the following pairs: visits 1-2: 0.80 and visits 3-4: 0.66 (Fig S2). The CV were 31% for visits 1-2 and 45% for visits 3-4. The estimated ICC for any of the other pairs of visits was not significantly different from 0 (Table S1). For saliva, we only found higher estimates of ICC for the pair visits 2 −3: 0.82 (Table S1). The estimated ICC for the remaining pairs was not significantly different from 0 (Table S1). Please see Supplementary Results (Table S1 and Figure S2) for correlations between baseline concentrations of oxytocin for each pair of visits of dataset A.

#### Dataset B

We estimated the ICC to be 0.49 for single measurements of baseline plasmatic oxytocin across the two visits. The mean CV was 42%. The number of measures that would have been required to achieve good reliability of a putative averaged measure was estimated to be 4.16 (Table 1).

### Effects of intranasal and intravenous oxytocin administration on salivary and plasmatic oxytocin concentrations

For salivary oxytocin, we found a significant treatment x time interaction (F(3, 122) = 18.29, p < 0.001). Post-hoc analyses revealed that the administration of oxytocin either by intranasal spray or nebuliser, but not the administration of intravenous oxytocin or placebo, resulted in significant increases of salivary oxytocin levels from baseline (<u>Baseline vs Post-administration:</u> Spray – t(122) = 7.06, p < 0.001; Nebuliser - t(122) = 7.61, p < 0.001; Intravenous - t(122) = 0.07, p = 0.99; Placebo - t(122) = 0.15, p = 0.99) (Fig. 3).

**Fig. 3.**
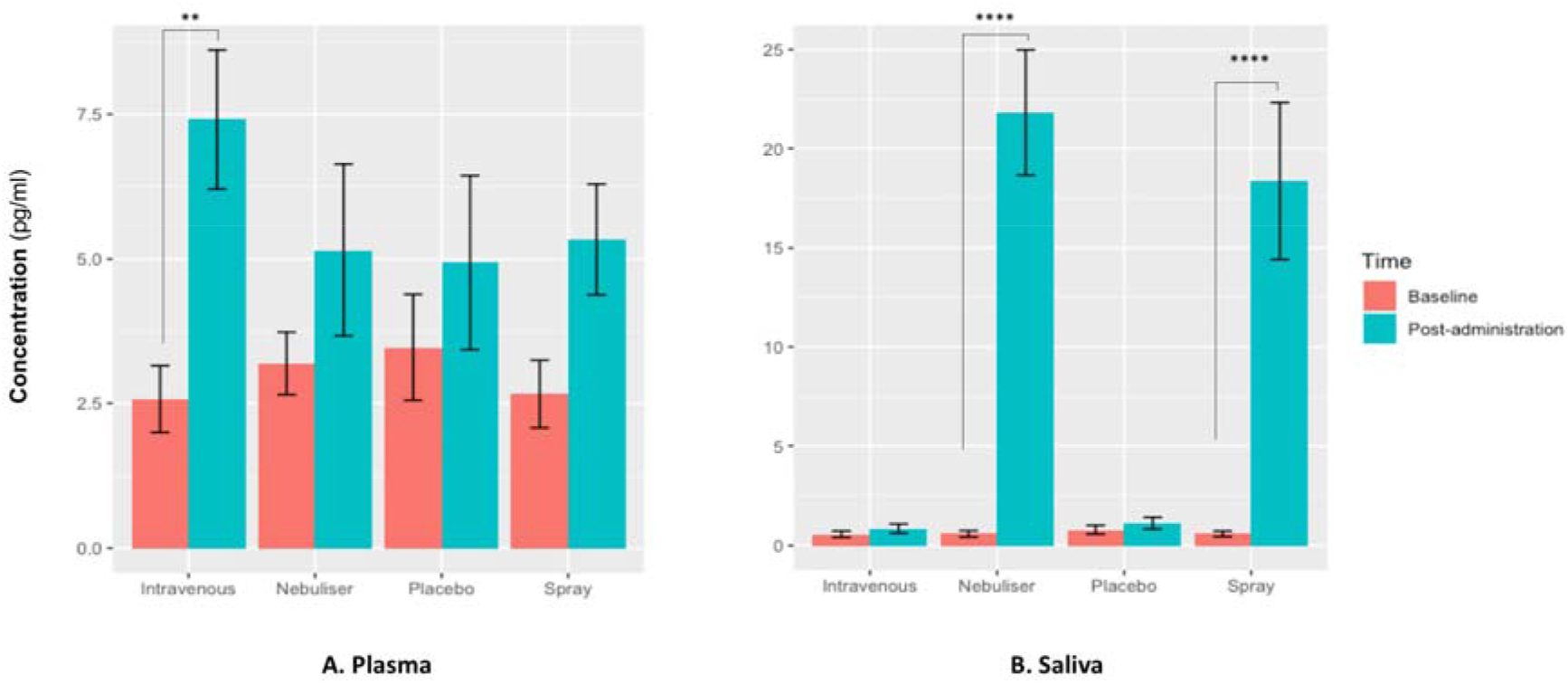
Effects of the administration of intranasal and intravenous oxytocin on salivary (A) and plasmatic (B) oxytocin. We examined the effects of treatment, time and treatment x time on salivary and plasmatic oxytocin in a 2-way analysis of variance. Post-administration samples were collected at 115 mins post-dosing. Statistical significance was set to p<0.05. **p = 0.001 and ****p<0.001, using Tukey for multiple testing correction during post-hoc investigation of significant interaction effects. Please note that while all the statistical analyses were conducted on log-transformed oxytocin concentrations, here we plot the raw values to facilitate interpretation.

For plasmatic oxytocin, we found a significant time x treatment interaction (F(3, 123) = 3.99, p = 0.02). Post-hoc investigations revealed that the intravenous administration of oxytocin resulted in a significant increase in plasmatic oxytocin, but placebo or intranasal administration of oxytocin using either the spray or the nebuliser did not produce any changes from baseline at this time-point (<u>Baseline vs Post-administration:</u> Spray – t(123) = 1.38, p = 0.52; Nebuliser - t(123) = 0.25, p = 0.99; Intravenous - t(123) = 3.73, p = 0.001; Placebo - t(123) = 1.54, p = 0.41) (Fig. 3).

### Association between salivary and plasmatic oxytocin at baseline and after administration of exogenous oxytocin

We did not find a significant correlation between oxytocin concentrations measured in saliva and plasma at baseline (r = 0.10, p = 0.18, BF = 3.02) (Fig. 4) or following the administration of exogenous oxytocin (<u>spray:</u> r = −0.21, p = 0.43, BF = 2.43; <u>nebuliser:</u> r = 0.07, p = 0.65, BF = 3.32; <u>intravenous:</u> r = −0.05, p = 0.84, BF = 3.27; <u>placebo:</u> r = −0.20, p = 0.45, BF = 2.48) (Fig. 5).

**Fig. 4.**
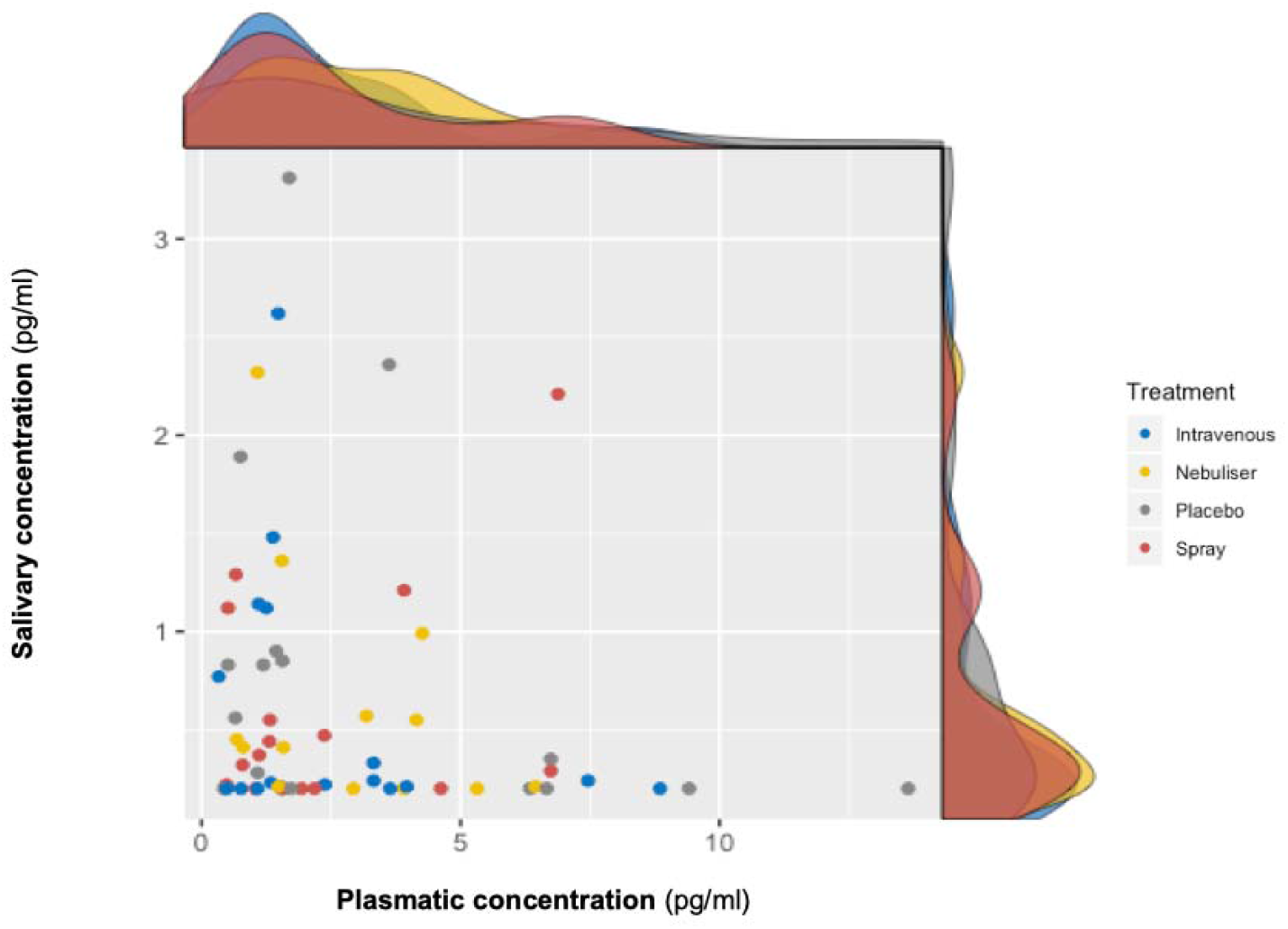
Association between salivary and plasmatic oxytocin concentrations at baseline. In this scatter plot, we depict the lack of association between salivary and plasmatic oxytocin concentrations at baseline (before any treatment administration). The density plots on the top of each axis show the distribution of oxytocin concentrations for each treatment level. Please note that while all the statistical analyses were conducted on log-transformed oxytocin concentrations, here we plot the raw values to facilitate interpretation.

**Fig. 5.**
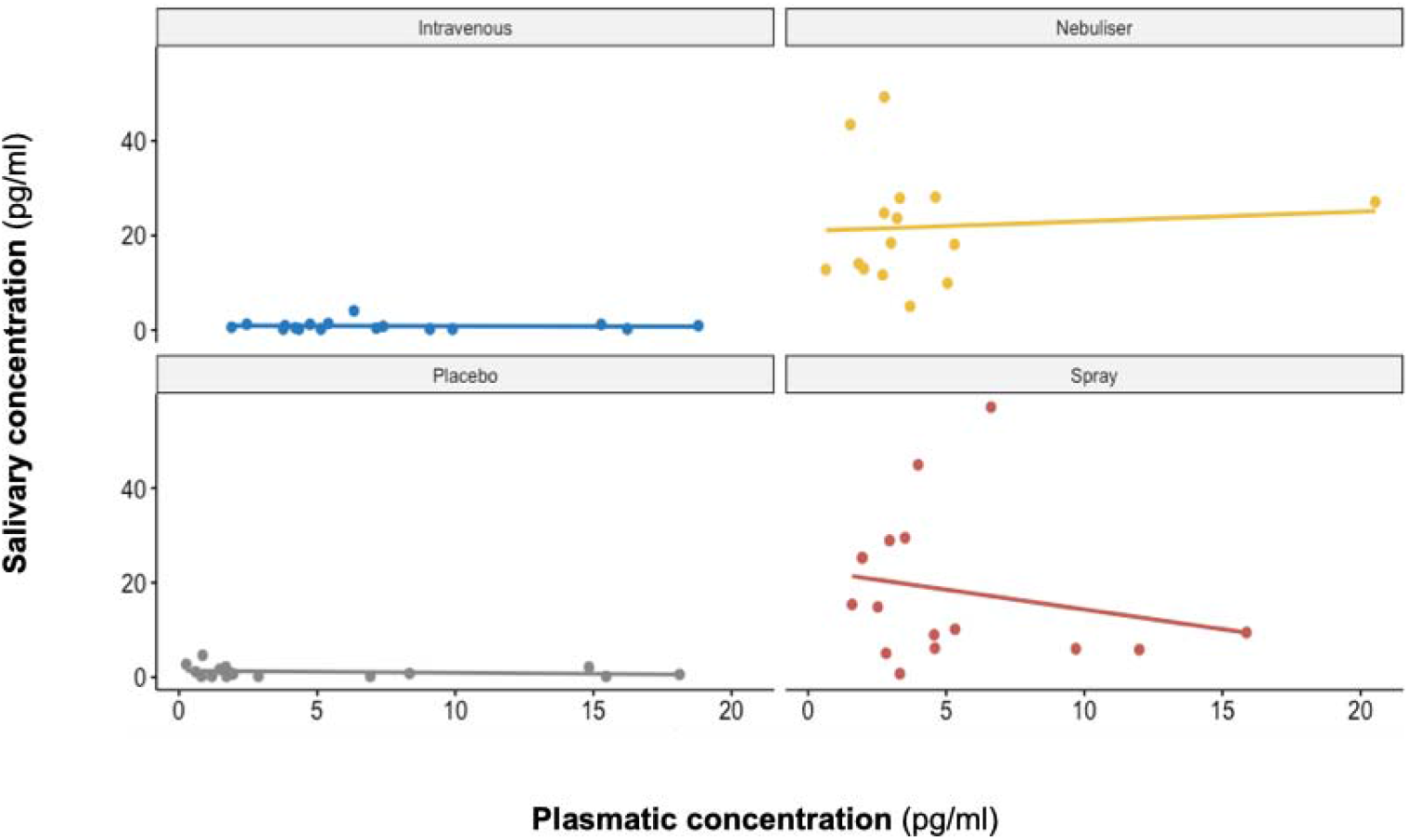
Association between salivary and plasmatic oxytocin after administration of intranasal and intravenous exogenous oxytocin. In these scatter plots, we depict the lack of association between salivary and plasmatic oxytocin concentrations at 115 mins after the administration of intravenous and intranasal oxytocin or placebo. Each panel depicts data from one out of the four treatment levels. Please note that while all the statistical analyses were conducted on log-transformed oxytocin concentrations, here we plot the raw values to facilitate interpretation.

## Discussion

Using radioimmunoassay, a gold-standard method for oxytocin quantification, we showed that i) single measures of baseline oxytocin concentrations in saliva and plasma are not stable within the same individual; ii) for plasma, we replicated this finding in an independent cohort; iii) intranasal administration of exogenous oxytocin increases salivary oxytocin, but intravenous administration of a considerable dose does not produce any changes; iv) salivary and plasmatic oxytocin do not correlate with each other either at baseline or after administration of exogenous oxytocin. We discuss our main findings below.

We reported poor reliability indexes for single measurements of baseline oxytocin in both plasma and saliva. This suggests that single measures of baseline oxytocin in either compartment cannot be consistently measured for the same individual. The reliability estimates found in the current study are in the range of those already described for other hypothalamic hormones such as vasopressin(32) or prolactin(33). We note that the mean CV for baseline concentrations of oxytocin in saliva and plasma is higher than four times the intra-assay variability of the radioimmunoassay we used (<10%). This suggests that the low reliability of endogenous oxytocin measurements across visits in the current study results from true intrinsic individual biological variability and not technical variability/error in the method used for oxytocin quantification.

While we made efforts to minimize variability in the conditions in which biological samples were collected across sessions in both datasets (i.e. asking participants to abstain from exercise the day before study; controlling the time of sample collection), several additional factors might underlie the intra-individual biological variability in plasmatic and salivary oxytocin we report here. These include differences across participants in sexual activity in the days preceding each visit, perceived stress, the amount of social interactions, or other non-acknowledged biological rhythms conditioning variations in oxytocin secretion throughout time(34, 35). For instance, one study showed that oxytocin is secreted in a pulsatile manner in healthy males even at rest(36). Therefore, sampling during different phases of the pulsatile release of oxytocin for plasma across sessions could explain the discrepancies in oxytocin measurements observed in the current study across visits. The time-interval between measurements do not seem to significantly impact on the reliability of baseline oxytocin. Although higher reliabilities could be identified between some consecutive pairs of sessions (separated by an average of about 6 days from each other: sessions 1-2 and sessions 3-4 for plasma measurements), other consecutive pairs of sessions (for instance, sessions 2-3) did not produce significant reliability estimates. If there are specific reasons explaining the higher reliability indices observed for the specific pairs of sessions, these reasons remain to be elucidated.

Our observation of poor reliability for single measurements of plasma and saliva oxytocin raises questions about the interpretation of previous evidence seeking to associate single measurements of baseline oxytocin with individual differences in a range of neuro-behavioural or clinical traits. Almost all these previous studies have relied on the collection of single samples from each individual. Our findings suggest measures of oxytocin could be inconsistent and thus it is unlikely that a single sample may accurately represent oxytocin physiology and thus capture relevant inter-individual differences. Reliability of measurements also impacts power to detect correlations against these measurements(37). To illustrate this point, in Supplementary Fig. S3 we provide the results of a set of simulations investigating how different reliabilities of oxytocin measurements impact the sample sizes necessary to detect significant correlations between oxytocin concentrations in peripheral fluids and neurobehavioral outcomes of different effect sizes. Given the less-than-perfect reliability of oxytocin measurements (ICC around 0.30) we show here, to detect the small-to-medium (r = 0.20 – 0.50) correlations typically reported in studies using these measurements(38), researchers would need sample sizes between 102 - 651 participants to reach minimally acceptable power (80%). Sample sizes in association studies of endogenous oxytocin measurements are typically below 100 participants(38).

Turning to our second main finding, salivary and plasmatic oxytocin did not correlate at baseline or after administration of exogenous oxytocin (irrespective of route), replicating previous observations of a null association between measurements in these two compartments(39). This finding was further supported through our Bayesian analysis which demonstrated that given our data the null hypothesis was about three times more likely than the alternative hypothesis. Two hypotheses could account for the lack of correlation between plasmatic and salivary oxytocin. First, as in all other studies in the field, we did not control or manipulate the rate of saliva flow in the current study. Lipid insoluble molecules, such as oxytocin, enter into saliva mainly via the tight junctions between acinar cells through ultrafiltration(40). If oxytocin does reach the saliva compartment through an ultrafiltration mechanism (which depends on saliva flow rate(9)), then it is possible that when saliva flow is stimulated, oxytocin measurements may better index its plasmatic concentrations. Second, we cannot discard differences in oxytocin degradation rates between saliva and plasma. Future studies should investigate these hypotheses further.

Studies have been using increases in salivary oxytocin after the intranasal administration of exogenous oxytocin to index systemic absorption and establish putative time-windows during which neurobehavioral effects of oxytocin administration may be expected. If oxytocin increases after intranasal administration reflected systemic absorption and transport from the blood, then we would have expected that intravenous oxytocin would have also increased salivary oxytocin. Moreover, we would also have expected that increases in plasmatic and salivary oxytocin after intranasal administration would reflect the typical ratio of their concentration as observed during baseline (with lower concentrations in saliva than in plasma). Our findings were not consistent with these expectations, supporting to the notion that increases in salivary oxytocin after its intranasal administration most likely reflect drip-down oxytocin from the nasal cavity. If this is the case, then oxytocin elevations in saliva are mainly driven by non-absorbed exogenous oxytocin and therefore their use to estimate levels of systemic absorption or predict treatment effects after intranasal oxytocin administration is not valid.

One may argue the absence of significant elevations of oxytocin in saliva after its intravenous administration may be explained by significant differences in the kinetics of oxytocin concentration variation between these two compartments. This may include a significant delay in the elevation of oxytocin in saliva after the beginning of its increase in the plasma (explaining why saliva and plasma concentrations of oxytocin are not correlated after its exogenous administration). While this may be possible, in a previous companion paper we showed that the peak in plasmatic oxytocin occurs immediately after the end of its intravenous administration – 115 min before our post-administration saliva sample collection(16). It is therefore unlikely that this large interval of time would not have captured a significant delay in saliva elevation of oxytocin if it really existed, especially when after this time-interval plasma oxytocin still remained elevated, compared to baseline levels.

A strength of our study is the replication of poor reliability for baseline plasmatic oxytocin in a second independent dataset, which strengths our confidence in the robustness of our reliability findings. However, we acknowledge the following limitations. First, we only considered baseline measures in our reliability analyses. As for other hypothalamic-pituitary-adrenocortical markers where evoked measures present higher reliability than baseline measures(41), stimuli-evoked release of endogenous oxytocin (i.e. after social interaction, stress, pain) might also present higher reliability. Our conclusions are also restricted to male participants and to the radioimmunoassay method of oxytocin measurement, precluding extrapolations to female populations or to other quantification methods. Also, due to time and logistical constrains during our MRI setup, we could only sample saliva before administration and at the end of the scanning period, leaving our analyses on the effects of the administration of exogenous oxytocin on its concentration in saliva restricted to one single-time point post-administration. It is possible that we may have missed peak increases in saliva oxytocin after the intravenous administration of exogenous oxytocin if they occurred between treatment administration and post-administration sampling.

In summary, single measurements of baseline levels of endogenous oxytocin in saliva and plasma are not stable and therefore their validity as biomarkers of the physiology of the oxytocin system is questionable. Salivary oxytocin is a weak surrogate for plasmatic oxytocin; hence, salivary and plasmatic oxytocin should not be used interchangeably. Finally, increases in salivary oxytocin after the intranasal administration of exogenous oxytocin most likely represent drip-down transport from the nasal to the oral cavity and not systemic absorption. Therefore, increases in salivary oxytocin after intranasal oxytocin administration should not be used to predict treatment effects.

## Acknowledgments

We would like to thank Sofia Vasilakopoulou and Jack Loveridge for their assistance in data collection. We also would like to thank Rosa Oliveira and Silia Vitoratou for their advice on statistical analysis. Most importantly, we thank all participants volunteering to both studies.

This study was part-funded by: an Economic and Social Research Council Grant (ES/K009400/1) to YP; scanning time support by the National Institute for Health Research (NIHR) Biomedical Research Centre at South London and Maudsley NHS Foundation Trust and King’s College London to YP; an unrestricted research grant by PARI GmbH to YP. Data collection for dataset B was supported by an IoPPN-MRC Excellence Studentship awarded to AG. MAM is in part supported by the National Institute for Health Research (NIHR) Biomedical Research Centre at South London and Maudsley NHS Foundation Trust and King’s College London.

## Author contributions

YP designed the study for dataset A; MM and AG designed the study for dataset B; YP collected the data for dataset A; AG for dataset B; DM analysed and interpreted the data; DM and YP wrote the first draft of the manuscript; MM and AG contributed critical revisions.

