## Supplementary for "Are single peripheral measurements of baseline oxytocin in saliva and plasma reliable biomarkers of the physiology of the oxytocin system in humans?"

### Methods

### Correlations between baseline oxytocin in saliva and plasma for each pair of visits from dataset A

### Correlations are often used as an index for reliability, even though they cannot provide information about the absolute agreement of two sets of measurements(1). Hence, to facilitate comparisons with previous reports, we also calculated Pearson’s correlation coefficients, with bootstrapping (1000 samples), to evaluate correlations between baseline concentrations of oxytocin for each pair of visits. The results of this analysis are presented below in the Table S1 and Figure S2.

### Bayesian analyses

### For the bayesian correlations, we used beta priors’ distributions centred around zero, with a width parameter of 1. An increase in Bayes Factor (BF) in our analyses corresponds to an increase in evidence in favour of the null hypothesis. To interpret BF, we used the Lee and Wagenmakers’ classification scheme(2): BF < 1/10, strong evidence for alternative hypothesis; 1/10<BF<1/3, moderate evidence for alternative hypothesis; 1/3 < BF < 1, anecdotal evidence for alternative hypothesis; BF > 1, anecdotal evidence for the null hypothesis; 3<BF<10, moderate evidence for the null hypothesis; BF > 10, strong evidence for the null hypothesis.

### Results

### Fig. S1 - Mean plasmatic and salivary concentrations of oxytocin across visits. We tested for differences in mean baseline plasmatic and salivary oxytocin across the four visits of dataset A in a repeated measures one-way analysis of variance. In dataset B, we tested for differences in mean baseline plasmatic oxytocin across the two visits using a paired T-test. Statistical significance was set to p< 0.05 (two-tailed). Box plots and violin plots depicting oxytocin concentrations for each visit; middle horizontal lines represent the median; boxes indicate the 25^th^ and 75^th^ percentiles.

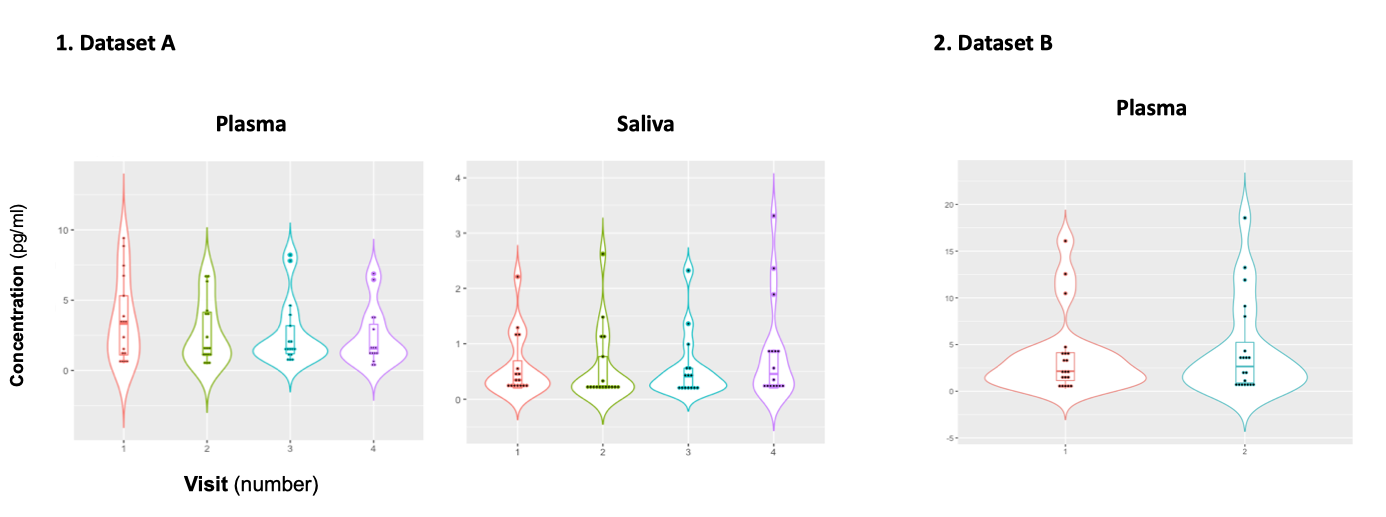

### Table S1 – Absolute and relative reliability of oxytocin measurements in plasma and saliva for each pair of the four visits included in study A. ICC – Intraclass correlation coefficient; CV – coefficient of variation; CI – confidence interval; SD – Standard Deviation; *H0:ICC is not significantly different from 0; statistical significance was set to p< 0.05 (two-tailed)).

|  |  |  | ICC | | | | | CV  Mean (SD) | Pearson r  r (p-value) |
| --- | --- | --- | --- | --- | --- | --- | --- | --- | --- |
|  |  |  |  | 95% CI | | F test* | p-value |  |  |
|  |  |  |  | Lower | Upper |  |  |  |  |
| Visit 1 vs 2 | Plasma | Single | 0.80 | 0.51 | 0.93 | F(15.15) = 8.45 | <1.00x10^-3^ | 31% (20) | 0.79 (p<0.001) |
|  | Saliva | Single | 0.11 | -0.51 | 0.62 | F(12.12) = 1.23 | 0.36 | 39% (38) | 0.11 (0.73) |
| Visit 1 vs 3 | Plasma | Single | 0.25 | -0.28 | 0.66 | F(15.15) = 1.64 | 0.18 | 43% (36) | 0.24 (0.36) |
|  | Saliva | Single | 0.08 | -0.54 | 0.60 | F(12.2) = 1.15 | 0.41 | 41% (33) | 0.07 (0.82) |
| Visit 1 vs 4 | Plasma | Single | 0.08 | -0.44 | 0.55 | F(15.15) = 1.17 | 0.51 | 55% (39) | 0.09 (0.75) |
|  | Saliva | Single | -0.17 | -0.67 | 0.42 | F(12.12) = 0.72 | 0.71 | 57% (45) | -0.18 (0.55) |
| Visit 2 vs 3 | Plasma | Single | 0.09 | -0.43 | 0.56 | F(15.15) = 1.19 | 0.37 | 51% (38) | 0.09 (0.75) |
|  | Saliva | Single | 0.82 | 0.52 | 0.94 | F(12.12) = 9.82 | <1.00x10^-3^ | 31% (31) | 0.83 (p<0.001) |
| Visit 2 vs 4 | Plasma | Single | -7.00x10^-3^ | -0.50 | 0.48 | F(15.15) = 0.99 | 0.51 | 57% (38) | -0.01 (0.98) |
|  | Saliva | Single | 0.35 | -0.21 | 0.74 | F(12.12) = 2.09 | 0.11 | 47% (39) | 0.38 (0.21) |
| Visit 3 vs 4 | Plasma | Single | 0.66 | 0.26 | 0.87 | F(15.15) = 4.69 | 2.00x10^-3^ | 45% (30) | 0.76 (p<0.001) |
|  | Saliva | Single | 0.30 | -0.24 | 0.71 | F(12.12) = 1.90 | 0.14 | 47% (41) | 0.36 (0.23) |

### Fig. S2 - Correlation between baseline oxytocin concentrations in plasma (left) and saliva (right) for each pair of the four sessions included in Study A. Colour grading represents Pearson’s coefficient of correlation (r), with bootstrapping 1000 samples. Statistical significance was set to p< 0.05 (two-tailed). Black crosses identify correlations that did not reach significance.

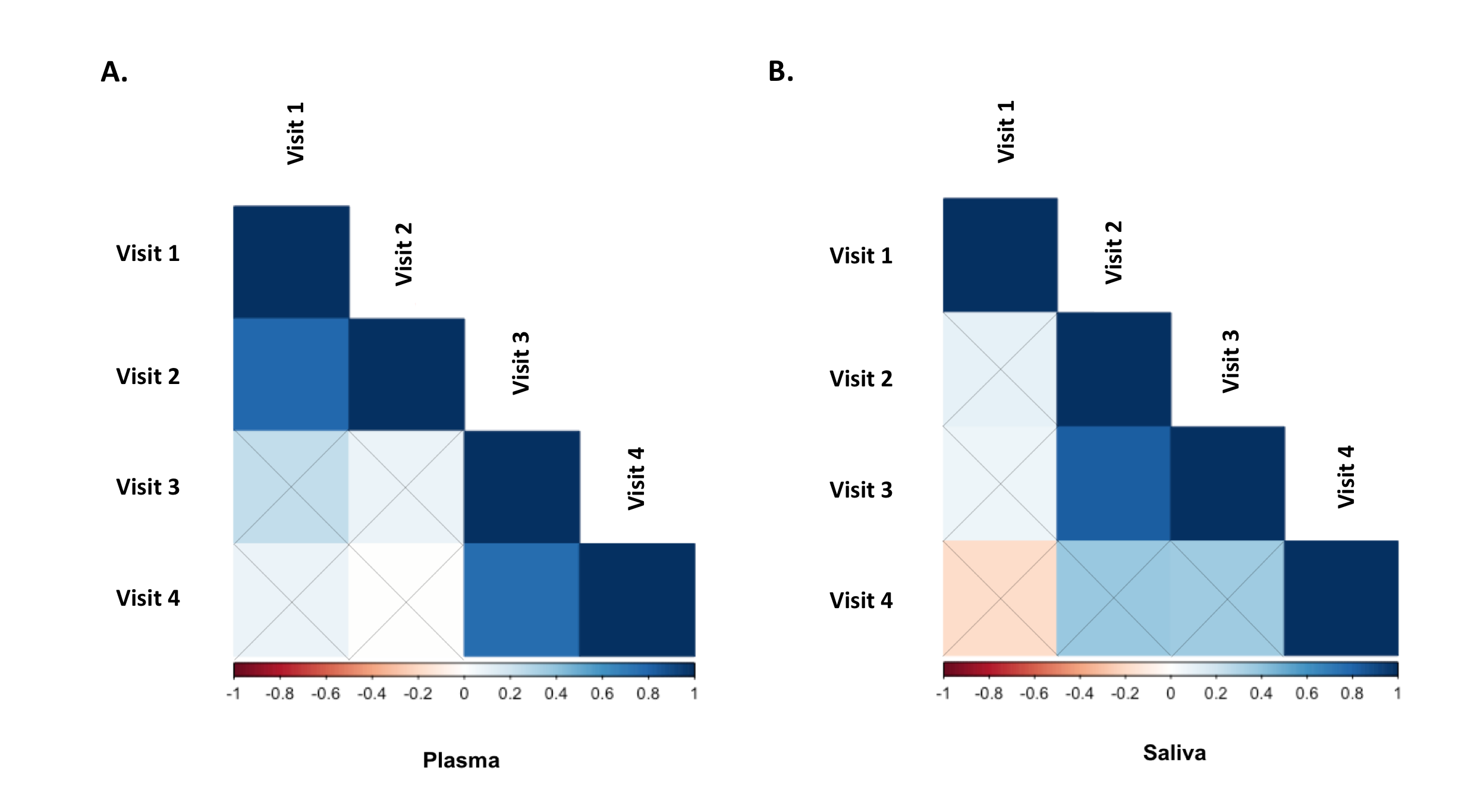

### Fig. S3 – Influence of less-than-perfect reliability of oxytocin measurements on the sample sizes required to detect significant endogenous oxytocin-outcome associations in neurobehavioral human oxytocin research. In this figure, we show the results of a set of simulations illustrating the impact less-than-perfect reliabilities of endogenous oxytocin measurements in peripheral fluids might have on the sample size required to detect significant oxytocin (A) – outcome (B) correlations of varying effect sizes in research studies. We conduct calculations for a minimally acceptable statistical power of 80% in a two-tailed parametric test. The outcome measure (B) was assumed to present perfect reliability. The figure was generated using the “pwr.r.test” function of the “pwr” R package. We specified “r” according to the attenuation formula below(3). ICC -. Intraclass correlation coefficient; r – Pearson’s correlation coefficient.

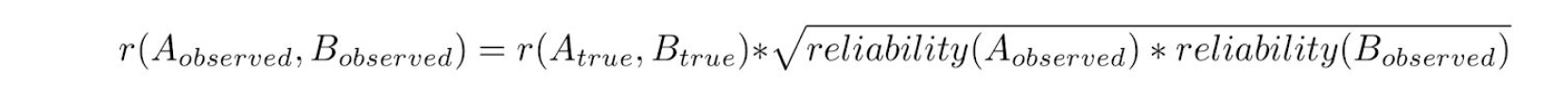

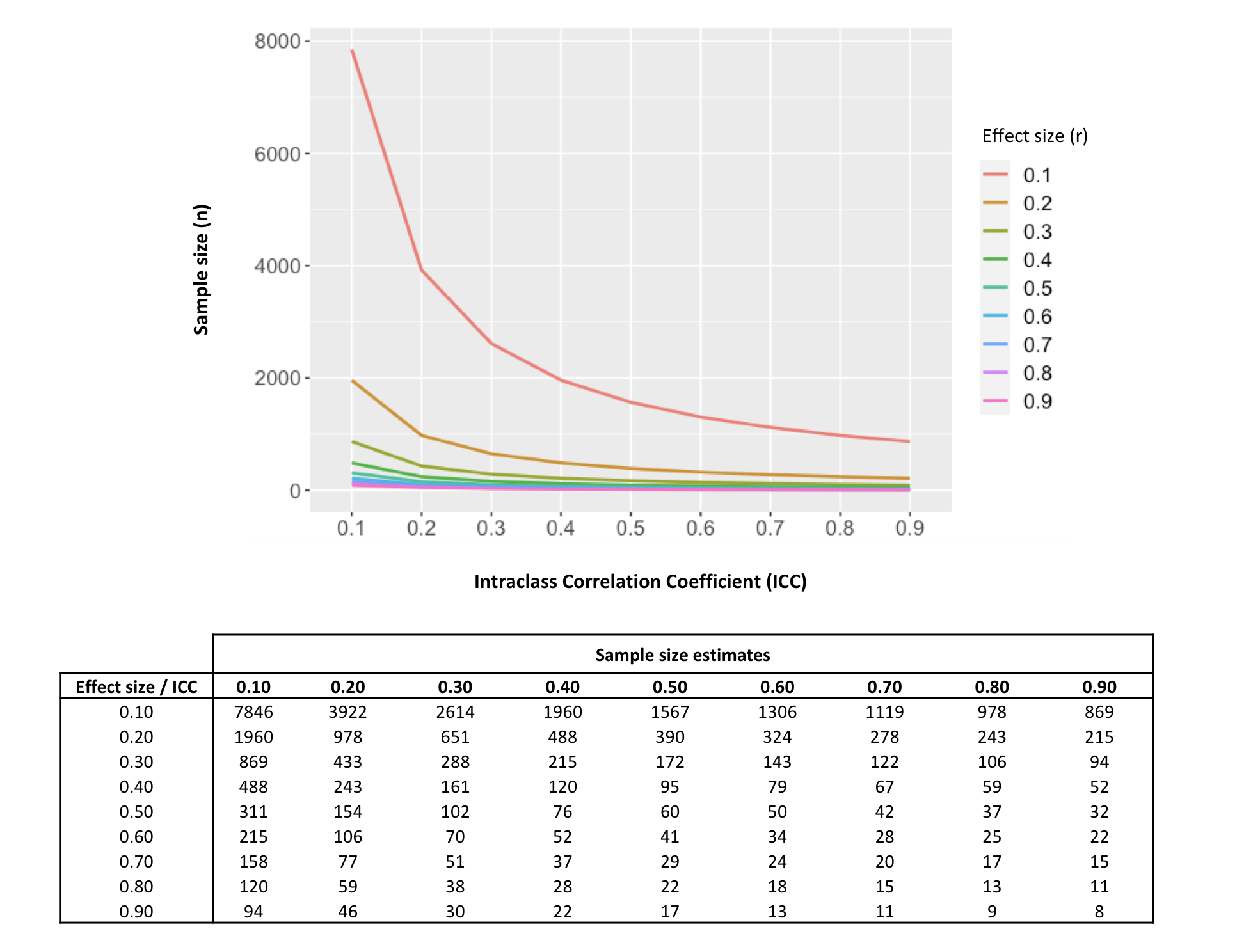
